# IL-33 regulates age-dependency of long-term immune dysfunction induced by sepsis

**DOI:** 10.1101/2022.01.15.476447

**Authors:** David F. Colon, Carlos W. Wanderley, Walter M. Turato, Vanessa F. Borges, Marcelo Franchin, Fernanda V. S. Castanheira, Daniele Nascimento, Douglas Prado, Mikhael Haruo Fernandes de Lima, Leila C Volpon, Silvia K. Kavaguti, Fernando Ramalho, Ana P. Carlotti, Fabio Carmona, Bernardo S Franklin, Jose C. Alves-Filho, Fernando Q. Cunha

## Abstract

Sepsis survival in adults is commonly followed by immunosuppression and increased susceptibility to secondary infections. However, the long-term immune consequences of pediatric sepsis are unknown. Here, we compared the frequency of Tregs, the activation of the IL-33/ILC2s axis in M2 macrophages, and the DNA methylation of epithelial lung cells from post-septic infant and adult mice. In contrast to adults, infant mice were resistant to secondary infection and did not show impairment in tumour controls upon melanoma challenge. Mechanistically, increased IL-33 levels, Tregs expansion, and activation of ILC2s and M2-macrophages were observed in post-septic adults but not infant mice. Impaired IL-33 production in post-septic infant mice was associated with increased DNA-methylation on lung epithelial cells. Notably, IL-33 treatment boosted the expansion of Tregs and induced immunosuppression in infant mice. Clinically, adults but not pediatric post-septic patients exhibited higher counts of Tregs and sera IL-33 levels. Hence, we describe a crucial and age-dependent role for IL-33 in post-sepsis immunosuppression.

## Introduction

Sepsis is a life-threatening multi-organ dysfunction caused by a dysregulated host response to an infection^1^. Adults that survived a sepsis episode frequently experience long-term immunosuppression, which increases the likelihood of secondary infections by opportunistic pathogens or the development of cancer ^2, 3, 4, 5, 6^. Despite these serious consequences, little is known about the development of post-sepsis immunosuppression in children. Longitudinal studies analysing the outcomes of survivor pediatric sepsis patients demonstrated that they did not have immunosuppression markers nor impairment in the quality of life after hospital discharge ^7, 8, 9^.

In adults, the development of long-term post-sepsis immunosuppression is associated with increased production of anti-inflammatory mediators, such as IL-10, IL-4, or TGF-ß ^5, 10, 11^; immune system dysfunction ^5, 12^; epigenetic alterations ^13, 14^; and expansion of specific cellular populations, including regulatory T cells (Tregs), B cells and M2-like macrophage ^15, 16, 17, 18^. Indeed, the inhibition of the Tregs suppressive capacity or genetic ablation of f*oxp3* on T cells significantly reduced the mortality due to secondary infection in sepsis-surviving adult mice ^15, 19^. Although the main regulator of post-sepsis immunosuppression is undetermined, recently, the alarmin IL-33 emerged as an important “rheostat” for the development of post-septic immunosuppression in adults ^17^.

IL-33, the latest member of the IL-1 family, is released by non-hematopoietic cells such as lung epithelial cells during injury ^20, 21, 22^. Alveolar epithelial cells have been reported as the major cellular sources of IL-33 ^23, 24^. IL-33 plays an important role in Th2-associated immune responses. After binding to its receptor (ST2), IL-33 induces the production of the Th2-associated mediators IL-4, IL-5, IL-10, and IL-13 by Th2 lymphocytes, mast cells, type 2 innate lymphoid cells (ILC2) and eosinophils ^25, 26^. Moreover, IL-33 drives the polarization of alternative-activated macrophages (M2) ^27^. Remarkably, post-septic IL-33 induced expansion of Tregs depends on ILC2s/M2-macrophages ^17^. However, the role of IL-33/ILC2s/M2-macrophages/Tregs axis in sepsis-surviving children remains unclear. Here, we investigated the long-term immune consequences of sepsis in surviving infants vs adults. Our findings demonstrate that, compared to adults, young mice that survived sepsis do not develop immunosuppression. These findings were recapitulated in sepsis-surviving pediatric patients in whom no Tregs expansion or increase in serum IL-33 levels was observed compared to adults. This study reveals that the long-term immunosuppression after sepsis might be an unappreciated age-dependent phenomenon and suggest that adult and pediatric sepsis-surviving patients require a different treatment approach.

## Methods

### Animals

Infant (2 weeks old) and adult (6 weeks old) C57BL/6 mice (wild-type, WT) were obtained from the animal facility of the Ribeirao Preto Medical School of University of São Paulo, São Paulo - Brazil. The animals were housed under standard conditions and received water and food *ad libitum*. Mice were housed in barrier cages under controlled environmental conditions (12/12 h of light/dark cycle, 55% ± 5% humidity, 23^°^C).

### Patients

Peripheral blood samples were collected from 21 sepsis-surviving patients (12 children and 9 adults), who were prospectively enrolled in the study after hospital discharge from the Intensive Care Unit of a tertiary-care university hospital at Ribeirão Preto. All patients fulfilled clinical or laboratory criteria for sepsis ^28^. Seventeen healthy volunteers (7 children and 12 adults) were recruited as controls. Pediatric disease severity was evaluated by PRISM (Pediatric Risk of Mortality) score and organ dysfunction by PELOD (Pediatric Logistic Organ Dysfunction) score ^29, 30^. The exclusion criteria included active haematological malignancy or cancer, chronic treatment with steroids, transplantation, HIV infection or advanced cirrhosis.

### Ethics approval and consent to participate

Animal studies were reviewed and approved by the Ethics Committee on the Use of Animals (CEUA) of the Ribeirão Preto Medical School, University of Sao Paulo, under protocol number 169/2011. The care and treatment of the animals were based on the Guide for the Care and Use of Laboratory Animals ^31^. The study was also approved by the Human Subjects Institutional Committee of the Ribeirão Preto Medical School, University of Sao Paulo, under protocol number 4886/2009. Written informed consent was obtained from patients or their parents/guardians/caregivers before enrolment and a blood sample was drawn.

### Bacterial culture

The caecal content of an adult C57BL/6 mouse was isolated, filtered through sterile gauze and grown in Brain Heart Infusion (BHI) (BD Diagnostic Systems, Sparks, USA) for 5 days, 37°C. The bacteria grown in this culture were washed two times with PBS, lyophilized and frozen on aliquots. One vial of bacteria was thawed and grown in a BHI medium, 37°C for 20 hours before each experiment. After two rounds of wash to remove the culture medium, bacteria were resuspended on sterile saline 0.9% and the number of bacteria was assessed by absorbance at 600 nm using a spectrophotometer (Molecular Devices, Sunnyvale, USA). To prepare *Pseudomonas aeruginosa* suspension, a stock strain isolated in a tertiary-care university hospital of Ribeirão Preto were prepared following the same procedures for caecal bacterial suspension.

### Experimental design

Infant and adult mice were submitted to sepsis by the intraperitoneal inoculation with 2 × 10^8^ CFU/cavity or 4 × 10^8^ CFU/cavity of bacterial suspension, respectively. Survival curves were prepared from the data recorded daily and serum biomarkers for organ functions were assayed at regular intervals. To assess the long-term phase of sepsis, animals undergoing sepsis received an intraperitoneal injection of ertapenem sodium (Merck Research Laboratory), 30 mg/kg to adult mice and 15 mg/kg to infant mice, beginning 6 h after sepsis and then every 12 h up to day 3. The survival rate was recorded daily for 5 days. At the end of this period, further experiments were performed with the surviving-sepsis mice who were euthanized on day 15 after sepsis induction by ketamine/xylazine overdose (>100 mg/kg, s.c., União Quimica, BR) followed by cervical dislocation.

To address the long-term sepsis immuno-consequences, two “double-hit” sepsis models were used: airway bacterial infection and tumor challenge model. For the airway second hit model, infant and adult sepsis-surviving mice were infected on the day 15 or 30 after sepsis induction with a virulent clinical strain of *Pseudomonas aeruginosa* suspension (8 × 10^5^ or 2 × 10^6^ CFUs/40 μL, respectively). The survival rate was recorded daily for up to 10 days.

For a tumor challenge model, melanoma B16 (ATCC) cells lines were cultured in RPMI containing 10% FBS (v/v), penicillin (100 U/mL) and amphotericin B (2 μg/mL). Before use, cells with 70% to 80% of confluence were detached with trypsin-EDTA 0.25% and washed in PBS twice. Subsequently, infant and adult sepsis-surviving mice were subcutaneously inoculated on the day 15 after sepsis with the B16 Melanoma cell line (5 × 10^4^ cells/mice). The tumor growth was followed from the day 0 to the day 15 after tumor inoculation. Tumor volumes were calculated according to the formula: tumor volume (mm^3^) = L × W2/2, where L represents the major axis (largest cross-sectional diameter) of the tumor, and W represents the minor axis. On day 7 after tumor transplant, a tumor density was assessed by IVIS (Xennogen IVIS Spectrum In Vivo Imaging System). Furthermore, on the day 15 after tumor inoculation, animals were euthanized by ketamine/xylazine overdose (>100 mg/kg, s.c., União Quimica, BR) followed by cervical dislocation and the tumor microenvironment were assessed by FACS.

### Bacterial counts

Bacterial counts were determined 6 h and 1, 7 and 15 days after infection, as previously described ^32^. Briefly, peritoneal exudate and blood samples were collected, serially diluted, plated on Muller-Hinton agar dishes (Difco Laboratories), and incubated at 37°C for 18 h, and CFU/ml were recorded.

### Glutamate Oxaloacetate Transaminase (GOT) activity

Animals were euthanized 6 h and 1, 7 and 15 days after infection, and blood samples were collected to measure hepatic damage by assessing GOT activity levels. The assays were performed with a commercial kit (Labtest, Brazil).

### Cytokine assays

Cytokine concentrations were measured by ELISA, using antibodies from R&D Systems according to the manufacturer’s instructions. The optical density of the individual samples was measured at 450 nm using a spectrophotometer (Spectra Max-250, Molecular Devices, Sunnyvale, CA).

### Flow cytometry

Aliquots of cells homogenate (1 × 10^6^ cells per tube) were suspended in buffer containing 2% FCS in PBS. For surface staining, cells were incubated with specific antibodies to F4/80 (BM8, eBioscience), CD206 (mannose receptor C type 1, MR; MR5D3, AbD Serotec), CD4 (GK1.5, eBioscience; H129.19, BD Biosciences), CD4 (RPA-T4, BD Biosciences, for human), CD45 (30-F11, BD Biosciences), CD11b (M1/70, BioLegend), CD11c (N418, BioLegend; Hl3, BD Biosciences), T1/St2 (IL-33R, DJ8, MD Biosciences) Lin (145-2C11; RB6-8C5; RA3-6B2; Ter-119; M1/70, BioLegend), Sca-1 (D7, BioLegend) or the appropriate isotype controls plus FcBlocker for 30 min. For transcription factor staining, cells were first stained for surface antigens, then fixed and permeabilized with mouse Foxp3 Buffer Set (BD Biosciences), according to the manufacturer’s recommendations. Cells homogenate were then incubated with specific antibodies to FoxP3 (FJK-16 s, eBioscience) or FoxP3 (150D/E4, eBioscience, for human) for 45 min. For intracellular cytokines staining, cells were stimulated for 4 h with eBioscience™ Cell Stimulation Cocktail and a protein transport inhibitor-containing Brefeldin (1.5uL/mL StopGolgi; BD Biosciences). Cells were then washed in FACS buffer, fixed immediately in formaldehyde (final concentration 4%) for 20 min on ice, washed and re-suspended in NP40 0.4% for 4 min at room temperature. Cells were washed twice in FACS buffer and stained with an antibody specific for IL-13 (eBio13A, eBioscience) or the appropriate isotype controls. Cells were analysed by FACSCanto using FCS Express V3 (De Novo Software, Los Angeles, CA).

### Lung Alveolar Epithelial Cells and type 2 Innate Lymphoid Cells immunophenotyping

The lung tissue from infant and adult post-septic mice was digested with Liberase TL (0.2 mg / mL, Roche) and DNase I (0.5 mg / mL, Sigma) for 45 minutes at 37°C under rotation. Then, total lung tissue was marked with a lineage antibody (Lin, anti-mouse CD3ε, Ly-6G / Ly-6C, CD11b, CD45R / B220, TER-119 PE, Biolegend), anti-EpCAM for 10 minutes at temperature environment. The epithelial alveolar cells were characterized as Lin-EpCAM^+^. For ILC2s characterization, total lung tissue was marked with a lineage antibody (Lin, anti-mouse CD3ε, Ly-6G / Ly-6C, CD11b, CD45R / B220, TER-119 PE, Biolegend), anti-CD45 (BD Bioscience) and anti-Sca-1 (BD Bioscience). Data were collected with a FACS Canto II (BD Biosciences) and then were analyzed with FlowJo 10.6.1 software (Treestar).

### *In vitro* T cell differentiation

CD4^+^ T cells were purified from spleen and lymph nodes with anti-CD4 and anti-CD25 microbeads negative selection (Miltenyi Biotech). Isolated cells were activated with plate-bound anti-CD3 (1 μg/mL) and soluble anti-CD28 (1 μg/mL, both BD Biosciences) in the presence of Tregs polarizing cytokines. Tregs were polarized with 1 ng/mL of TGF-β1 (R&D Systems). Cells were cultured for 72 hours and collected for flow cytometry analysis.

### Macrophage differentiation and polarization

BMDM were differentiated as described previously ^27^. Bone marrow cells from infant and adult naïve mice were cultured in the presence of 20% of L929 cell culture supernatant (v/v) for 7 days. After differentiation, cells were seeded at a density of 1 × 10^6^ cells per well in 12-well plates and stimulated with LPS (100 ng/mL, M1-like - Sigma-Aldrich), IL4 (10 ng/mL, M2-like, R&D Systems) or medium (M0 macrophages). After 48 hours, the cells were used for FACS and supernatants for cytokine analysis by ELISA.

### Coculture of macrophages with T cells

Subsets of macrophages (M0, M1 or M2, 5 × 10^5^ per well) were co-cultured for 4 days with effector CD4^+^ T cells (CD4^+^ CD25−T cells, 5 × 10^5^ per well) purified from the spleen of naive infant and adult mice, in the presence of IL-2 (10 ng ml−1, R&D Systems), anti-IL-10 (50 μg ml−1, clone JES052A5, R&D Systems) and stimulated with polyclonal anti-CD3 (1 μg ml−1, BD Biosciences) in U-bottom 96-well plates. Cells were then stained for FoxP3 and CD4 and analyzed by FACS.

### Total DNA imprinting methylation

Lung Lin-cells of infant and adult sepsis-surviving animals were isolated and the total DNA was extracted using a commercial kit (Wizard® Genomic DNA Purification Kit, Promega) according to the manufacturer’s recommendations. Afterwards, the total DNA methylation signature was evaluated in 100 ng of DNA using the commercial kit (Imprint Methylated DNA Quantification kit, Sigma-Aldrich), according to the manufacturer’s recommendations.

### Gene expression by real-time PCR

Total RNA from the lung was extracted using TRIZOL reagent (Invitrogen) or RNeasy kit (Qiagen) according to the manufacturer’s instructions. Total RNA (2 μg from tissue and 1900 ng from cells) was reverse-transcribed using high-capacity cDNA RT Kit (Applied Biosystem). A High-Capacity cDNA kit (Life Technologies) was used and results were analyzed by quantitative RT-PCR with a Vii 7 Real-time PCR system (Applied Biosystems). The comparative threshold cycle method and internal control (Gapdh) was used for the normalization of the target genes. Real-time PCR was performed using the following primers: *Foxp3*, F- TTCTCCAGGACAGACACAACT / R- GTTGCTGTCTTTCCTGGGTGTA, *Ctla4*, F- TGTTGACACGGGACTGTACCT / R- CGGGCATGGTTCTGGATCA, *Tgfb1*, F- CCTGTCCAAACTAAGGC / R- GGTTTTCTCATAGATGGCG, *Gitr*, F- AAGGTTCAGAACGGAAGT / R- GGGTCTCCACAGTGGTAC, and *Gapdh*, F- GGGTGTGAACCACGAGAAAT / R- CCTTCCACAATGCCAAAGTT.

### Western blot analysis

Mice were terminally anesthetized and the lungs were collected. Samples were homogenized in a lysis buffer containing a mixture of proteinase inhibitors (Tris-HCl 50 mM, pH 7.4; NP-40 1%; Na-deoxycholate 0.25%; NaCl 150 mM; EDTA 1 mM; PMSF 1 mM; Aprotinin, leupeptin and pepstatin 1 μg/ml). Proteins were separated by SDS-polyacrylamide gel electrophoresis and trans-blotted onto nitrocellulose membranes (Amersham Pharmacia Biotech). The membranes were blocked with 5% dry milk and incubated overnight at 4°C with rabbit polyclonal antibody against p-Smad2/3 (1:200; ab272332, Abcam), Smad2/3 (1:200; ab217553, Abcam), p-CREB (1:200; ab32096, Abcam) and CREB (1:200; ab32515, Abcam). The membranes were incubated with a secondary antibody (Jackson ImmunoResearch). Immunodetection was performed using an enhanced chemiluminescence light-detecting kit (Amersham Pharmacia, Biotech) for 1 min. Optical densitometry was measured following normalization to the control using Scientific Imaging Systems (Image labTM 3.0 software, Biorad Laboratories, Hercules CA).

### Statistical analysis

The data (except for the survival curves) are reported as the mean ± standard error of the mean (SD) of values obtained from at least two independent experiments. The means of different treatments were compared by analysis of variance (ANOVA) followed by Bonferroni’s Student’s *t*-test for unpaired values. Bacterial counts were analyzed by the Mann-Whitney *U* test. The survival rate was expressed as the percentage of live animals, and the Mantel-Cox log-rank test was used to determine differences between survival curves. *P* <0.05 was considered significant. All statistical analyses were performed using GraphPad Prism version 9.00 (GraphPad Software, USA).

## Results

### Absence of post-sepsis-induced immunosuppression in infant mice

To assess the immune consequences of sepsis in infants, a “double-hit” sepsis model was used. For this, we induced sepsis in infant (2 weeks old) or adult (6 weeks old) mice by intraperitoneal (i.p) injection of bacterial suspension, as previously reported ^33^. Mice were treated with the antibiotic ertapenem, beginning 6 h after sepsis induction and then every 12 h up to day 3 (**Supplemental file 1: Figure S1A**). As expected, ertapenem significantly reduced the mortality rate of the animals and improved the progressive recovery of body weight (**Supplemental file 1: Figure S1B, C**). Concentrations of GOT (liver injury) in serum and bacteria loads in blood were also significantly reduced by ertapenem treatment (**Supplemental file 1: Figure S1D, E**). To address the immune status of sepsis-surviving mice, we administered a non-lethal dose of *P. aeruginosa*, a common post-sepsis opportunistic pathogen ^34, 35^. For that, infant and adult naïve mice were infected intranasally with different doses of *P. aeruginosa* suspension (10^5^ − 10^6^ CFU) and the survival rate was recorded. The doses of 8 × 10^5^ CFU/40 μl to infants and 2 × 10^6^ CFU/40 μl to adult mice that led to 80% of survival in both groups were selected to be used as the second infection hit (**Supplemental file 1: Figure S1F, G**). Further, sepsis-surviving mice were submitted to the second hit of *P. aeruginosa* on day 15 or 30 after sepsis induction. In agreement with previous findings ^15^, adult mice that survived sepsis become highly susceptible to secondary *P. aeruginosa* infection, indicating the development of post-sepsis immunosuppression. Notably, sepsis-surviving infant mice were resistant to a secondary *P. aeruginosa* infection (**Fig. 1A, B**) and displayed no changes in the lung bacterial load (**Fig. 1C**). The results suggest that post-septic infant mice do not develop immunosuppression. Post-sepsis immunosuppression has been described as a key risk factor for the development of lung carcinoma and melanoma growth ^16, 36^. Therefore, to confirm the post-sepsis immunosuppression, we injected surviving mice with the murine melanoma cell line (B16). Post-sepsis adult mice displayed an increase in the density and growth of implanted B16 tumors. Confirming the absence of immunosuppression in infant mice, B16-challenged infant mice did not show differences in the tumor growth when compared to the infant sham group (**Fig. 1D - F**). Additionally, infant post-septic spleen CD4^+^ T cells proliferation capacity was not compromised after *in vitro* stimulation with anti-CD3/anti-CD28 compared to adults **(Supplemental file 2: Figure S2**). Altogether, these results show that sepsis-surviving infant mice do not develop post-sepsis immunosuppression.

**FIGURE 1.**
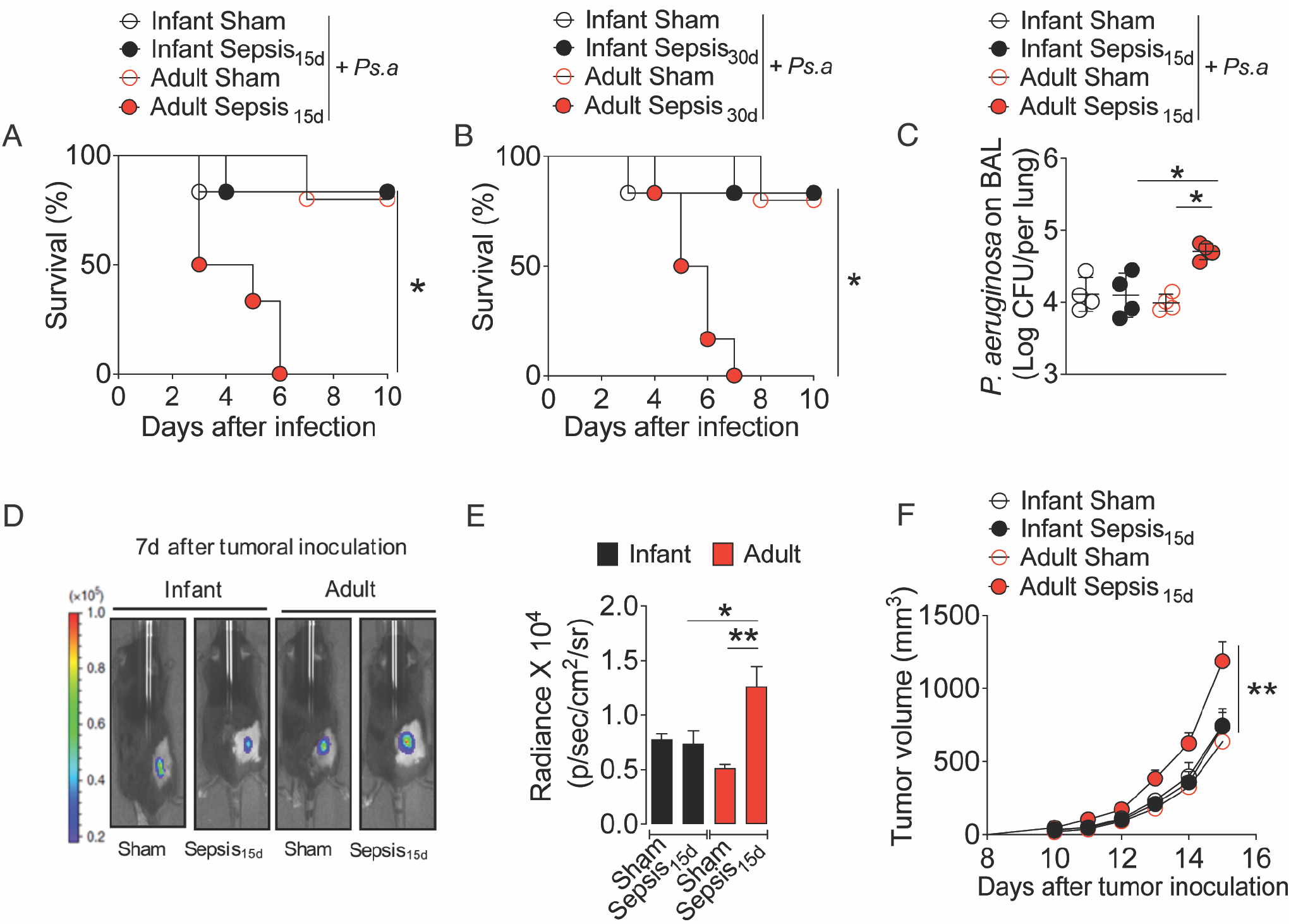
Absence of post-sepsis-induced immunosuppression in infant mice. Infant and adult mice were intraperitoneally injected with 2×10^8^ and 4×10^8^ colony-forming units (CFU) of cecum bacteria, respectively, and after six hours, the ertapenem antibiotic therapy (Abx, 15 mg/kg for infant and 30 mg/kg for adults) was initiated and maintained for 3 days, via intraperitoneal (i.p) twice a day. On days 15 **(A)** or 30 **(B)**, the surviving mice were submitted to the second hit with an intranasal injection of *P. aeruginosa*, 8 × 10^5^ CFU for infant and 2 × 10^6^ CFU for adults, and the survival percentage was calculated with the data recorded daily for 10 days. **(C)** The logarithm of CFU per lung (Log CFU/lung) was determined by seeding the bronchoalveolar lavage (BAL) collected 12 h after *P. aeruginosa* infection in the 15 days sepsis-surviving (Sepsis_15d_) mice. We also submitted the Sepsis_15d_ mice to B16 melanoma cells subcutaneous inoculation (5 × 10^4^ cells/mice) and, after 7 days, the tumor density **(D)** was evaluated by in vivo imaging system (IVIS) and the average radiance was expressed in 10^4^ photons per second per square centimeter per steradian (p/sec/cm2/sr) **(E)**. The tumor volume **(F)** in cubic millimeters (mm3) was also evaluated daily from day 10 to day 15 after tumoral cells challenging. Data are mean ± SD, n=6-8 per group and are representative of 2-3 independent experiments. * *p*<0.5 and ** *p*<0.01 (**A - B**, Mantel-Cox log-rank test; **C – F**, one way-ANOVA, Bonferroni’s).

### Lack of Tregs expansion in sepsis-surviving infant mice

Because Tregs are a major cellular driver of the post-sepsis immunosuppression in adults ^15, 16, 17, 37, 38^, we sought to assess the *in vivo* expansion of Tregs population in sepsis-surviving infant mice. Remarkably, in contrast to post-sepsis adults, post-septic infant mice displayed no increase in the spleen Tregs population, as well as no changes in *Foxp3* or *Tgfb1* expression on isolated spleen Tregs, conventional T cells (Tconv) and total spleen tissue (**Fig. 2A – D and Supplemental file 3: Figure S3A**). Concordantly, we found that the expansion in the frequency of both iTregs (induced regulatory T cells, CD4^+^Foxp3^+^Neuropilin1^−^) and nTregs (natural regulatory T cells, CD4^+^Foxp3^+^Neuropilin1^+^) did not occur in post-septic infant animals compared to adults (**Supplemental file 3: Figure S3B - D**). We also observed a significant reduction in the *ex vivo* proliferation capacity of both total CD4^+^ T spleen cells and Tregs from post-septic infant mice (**Supplemental file 3: Figure S3E, F**). To confirm these findings, we carried out the intratumoral Tregs frequency in post-septic bearing breast tumor mice. Consistent with the aforementioned findings, we observed no increase in the frequency of Tregs in the tumor microenvironment of post-septic infant mice compared to the adult counterparts (**Fig. 2E – F**). Moreover, compared to infant mice in post-septic adult animals we observed a reduced frequency of cell subtypes that are essential for the control of tumor growth (IFN-γ-producing CD4^+^ and CD8^+^ cells) ^39, 40, 41, 42^ (**Fig. 2G – H**). Corroborating this, post-septic adult mice presented a higher ratio of Tregs/ IFN-γ-producing CD8^+^ cells and Tregs/CD4 IFN^+^ cells, which indicates a more severely immunosuppressed tumor microenvironment when compared to the post-septic infant group (**Fig. 2I**). To investigate the mechanisms involved in the failure of Tregs expansion, we assessed whether CD4 T cells from infant mice could have impaired the Tregs differentiation. We stimulated the *in vitro* Tregs differentiation of CD4 T cells from adult and infant mice cultured with TGF-ß. We found that the differentiation to the regulatory profile (CD4^+^Foxp3^+^) was similar in both groups. No differences were found in the expression of hallmark genes associated with the Tregs profile (*Foxp3, Tgfb1, Gitr, Ctla4*, and *Tigit*) (**Supplemental file 4: Figure S4A, B**). Validating these findings, we observed that the early differentiation to the regulatory profile (CD4^+^CD25^hi^ T cells) was similar in both groups (**Supplemental file 4: Figure S4C**). We then assessed the early phosphorylation of SMAD2/3 and CREB, essential transcription factors involved in Tregs stability and differentiation ^43, 44^, after TGF-ß stimulation. TGF-ß increased the expression of activated SMAD2 (p-SMAD2) and CREB (p-CREB) in a time-dependent manner (**Supplemental file 4: Figure S4D**), indicating that Tregs differentiation is an age-independent process. Collectively, these results suggest that the reduced expansion of Tregs in sepsis-surviving infant mice is independent of the intrinsic fitness of the infant Tregs.

**FIGURE 2.**
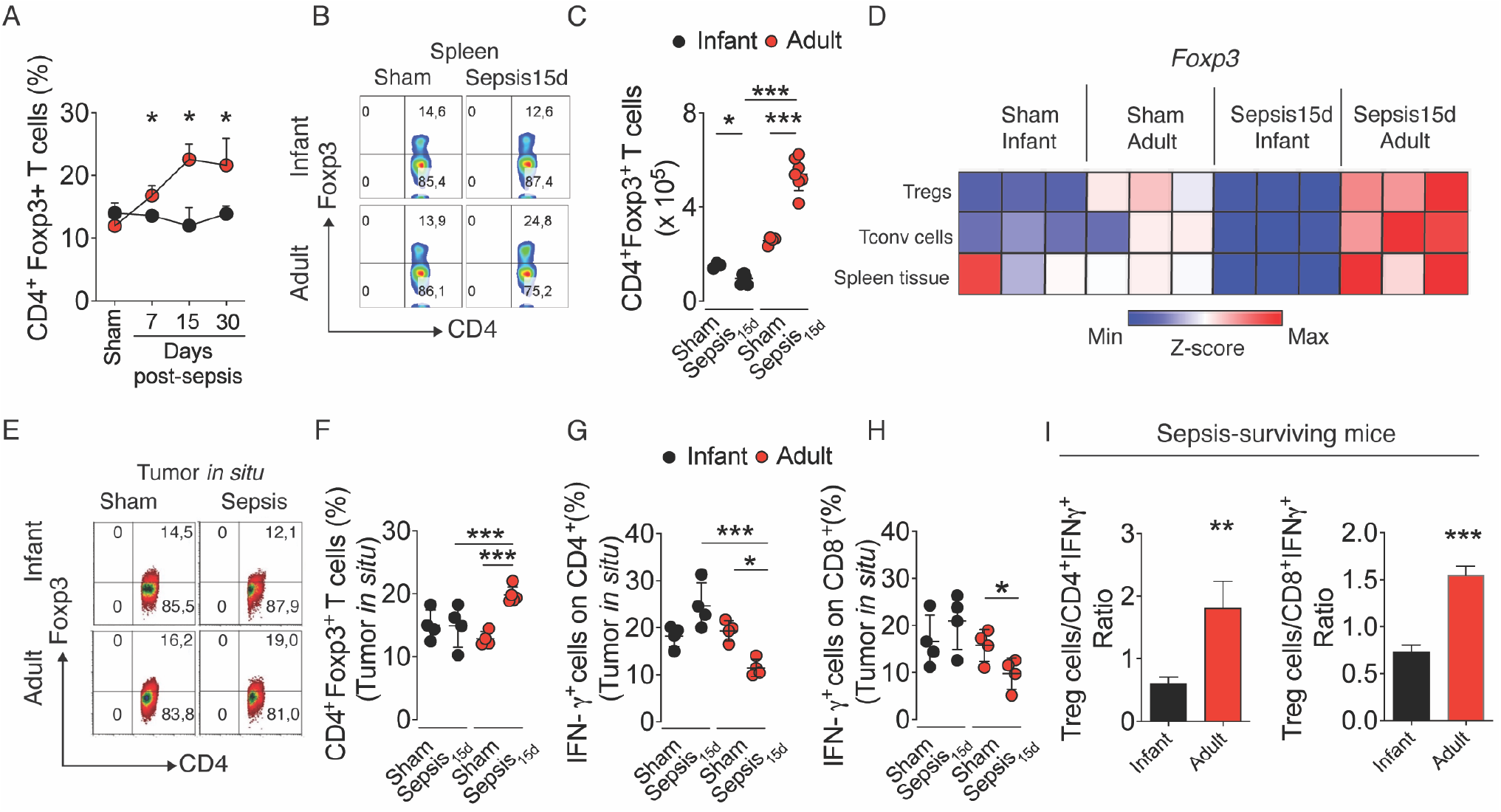
Lack of Treg expansion in sepsis-surviving infant mice. Infant and adult mice were intraperitoneally injected with 2×10^8^ and 4×10^8^ colony-forming units (CFU) of cecum bacteria, respectively, and after six hours, the ertapenem antibiotic therapy (Abx, 15 mg/kg for infant and 30 mg/kg for adults) was initiated and maintained for 3 days, via intraperitoneal (i.p) twice a day. The frequency in percentages of spleen T regulatory cells (Tregs) **(A)** on sepsis-surviving mice in the days indicated in the figure, as well as the representative flow cytometry plots **(B)** and the absolute number of Tregs in the 15 days sepsis-surviving (Sepsis_15d_) mice **(C)** addressed by flow cytometry. **(D)** Heat map of *Foxp3* expression in spleen Tregs (CD4^+^CD25^+^), conventional T cells (Tconv, CD4^+^CD25^−^) and whole spleen tissue from Sepsis_15d_ mice. Sepsis_15d_ mice were inoculated with B16 melanoma cells (5 × 10^4^ cells/mice) and, after 15 days, the tumor was removed and the intratumor cells were evaluated by flow cytometry. **(E)** Representative flow cytometry plots and **(F)** frequency of intratumoral Tregs. Frequency of **(G)** CD4^+^ and **(H)** CD8^+^ IFNγ-producing cells. The ratio between Tregs cells and CD4^+^ or CD8^+^ IFNγ-producing cells **(I)** in the tumor microenvironment from sepsis-surviving mice. Data are mean ± SD, n=6-8 per group and are representative of 2-3 independent experiments. * *p*<0.5 and **** *p*<0.0001 (**A** and **I**, *t-*test and **A, F – H** one way-ANOVA, Bonferroni’s; **D**, Z-score normalized heat map).

### Sepsis does not increase M2-like macrophages profile in post-septic infant mice

To further examine the mechanism of the dampened expansion of Tregs in post-septic infant mice *in vivo*, we assessed the effect of sepsis in infant M2-like macrophages polarization. The M2-like macrophages have a particularly important role in post-sepsis immunosuppression through the induction of Tregs differentiation in adults ^17, 45, 46, 47^. We found that whereas post-septic adult mice showed an increased frequency in peritoneal M2-like macrophages (F4/80^+^ CD206^+^) and reduced bacterial killing, the frequency of M2-like macrophages was significantly reduced in post-septic infant mice, with no impairment in the bacterial killing (**Fig. 3A, B**). Furthermore, the adult post-sepsis immunosuppression triggered the expression of M2 hallmark genes (*Ym1, Mrc1* and *Arg1*) in the peritoneal cells, total lung tissues, and alveolar macrophages. In contrast, no such increased expression was found in post-septic infant mice (**Fig. 3C – E**). Then, we assessed whether infant mice could have a cell-intrinsic impairment in the M2 polarization. For that, we compared the M2 polarization of BMDMs from adult and infant mice *in vitro*. Consistent with the Tregs findings, we found no impairment in the M2 polarization in infant mice (**Fig. 3F and Supplemental file 5: Figure S5A**). Moreover, hallmarkM2-associated genes (*Ym1, Mrc1* and *Arg1*) were similarly upregulated in a time-dependent manner in both infant and adult BMDMs in response to IL-4 (**Supplemental file 5: Figure S5B**). There were also no differences in the production of IGF-1, a tissue damage resolution mediator ^48^, as well as in CCL22, a Tregs attracting chemokine ^49^ between adult and infant mice (**Supplemental file 5: Figure S5C**). We then investigated whether M2 infant macrophages are less able to induce Tregs than the adult M2 macrophages. We cocultured M2-like macrophages from infant and adult mice with isolated CD4^+^ T cells from adult or infant mice, and the Tregs differentiation was assessed. Although adult and infant M2 macrophages were able to induce the Tregs differentiation, this was even more prominent in the presence of infant M2 macrophages (**Fig. 3G**) showing that there is no defect in the ability of M2 macrophages from infant mice to induce Tregs *in vitro*. Altogether, these findings suggest that events upstream to M2 macrophage polarization and Tregs expansion are involved in the dampened development of post-sepsis immunosuppression in infant mice.

**FIGURE 3.**
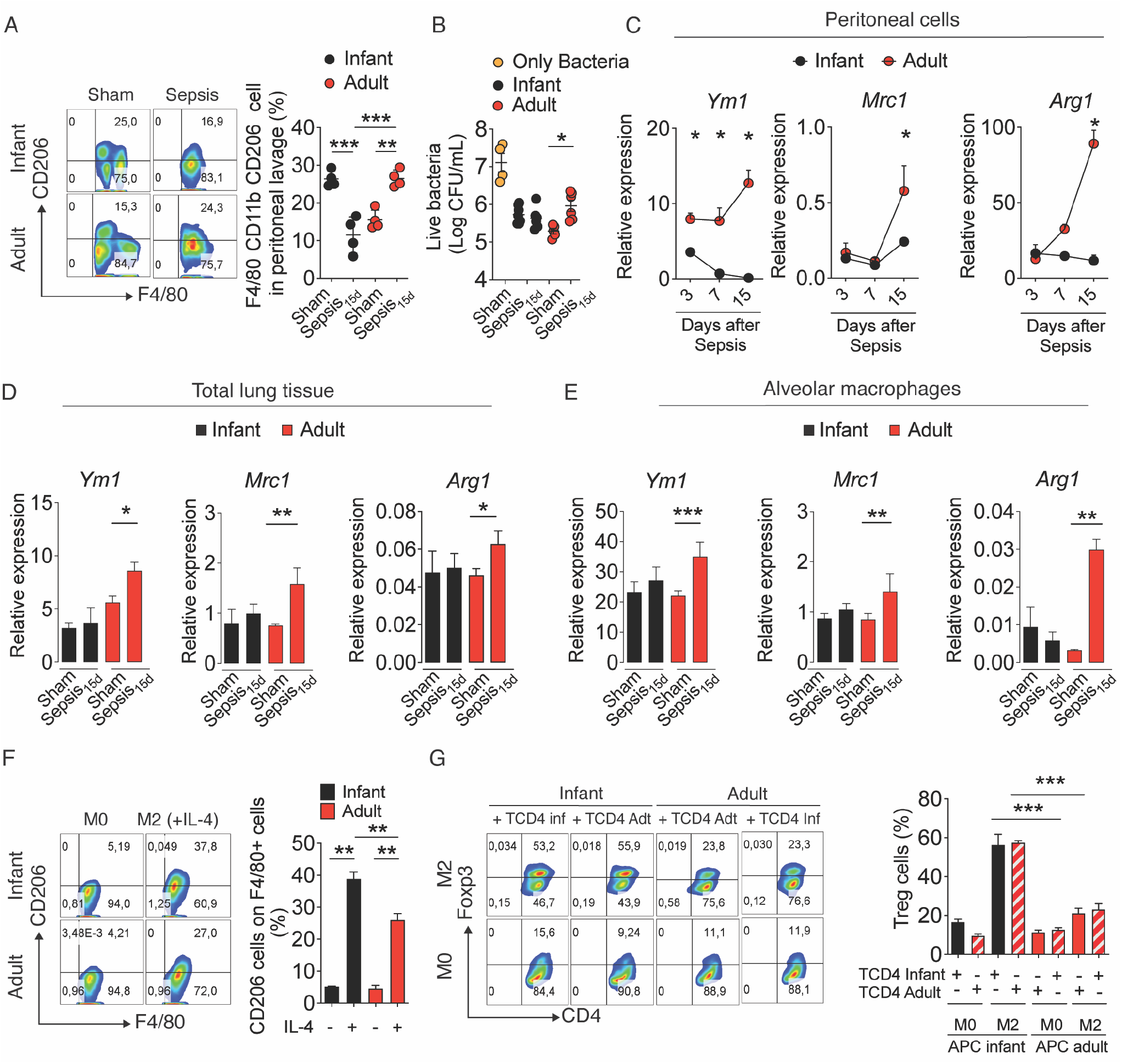
Sepsis does not increase the M2-like macrophages profile in post-septic infant mice. Infant and adult mice were intraperitoneally injected with 2×10^8^ and 4×10^8^ colony-forming units (CFU) of cecum bacteria, respectively, and after six hours, the ertapenem antibiotic therapy (Abx, 15 mg/kg for infant and 30 mg/kg for adults) was initiated and maintained for 3 days, via intraperitoneal (i.p) twice a day. **(A)** Representative flow cytometry plots and frequency in the percentage of M2-like macrophages (F4/80^+^CD206^+^) in the peritoneal exudates of 15 days sepsis-surviving (Sepsis_15d_) mice. **(B)** Live bacteria after killing assay of *E. coli* by peritoneal macrophages from Sepsis_15d_ mice. *Ym1, Mrc1* and *Arg1* relative expression in the days indicated in the figure or Sepsis_15d_ mice in the **(C)** peritoneal fluid exudates **(D)** total lung tissue, and **(E)** alveolar macrophages assessed by qPCR. Infant and adult bone marrow-derived macrophages (BMDMs) were polarized in the presence of IL-4 (10 ng/mL) for 48 h. **(F)** Representative Flow cytometry plots and frequency in the percentage of F4/80^+^CD206^+^ macrophages. **(G)** Bone marrow-derived macrophages (M0) or M2-like polarized macrophages (BMDM + IL-4, 48h) from infant or adult mice were co-cultured with infant or adult spleen-isolated CD4^+^CD25^−^ T cells in the presence of anti-εCD3 (1μg/mL) and the Tregs differentiation was addressed 72 h later by flow cytometry and shown as representative flow cytometry plots and frequency in the percentage of CD4^+^Foxp3^+^ T cells **(G)**. Data are mean ± SD, n=4-6, representative of two experiments, \**p*<0.05, \*\**p*<0.01, *** *p*<0.001 and **** *p*<0.0001 (one way-ANOVA, Bonferroni’s; **C**, *t-*test).

### Sepsis does not increase the ILC2/IL-33 axis in sepsis-surviving infant mice

Recently, the alarmin IL-33, a member of the interleukin (IL)-1 family, has been identified as a major player in the post-sepsis immunosuppression by activating the Tregs/M2-like macrophages axis ^17^. We, therefore explored the effect of IL-33 administration on Tregs/M2-like macrophages axis in infant mice. IL-33 administration resulted in a robust expansion of peritoneal M2-like macrophages and spleen Tregs in both infant and adult mice in an IL-33 receptor (ST2)-dependent manner (**Fig. 4A, B**). Noticeably, compared to the adults, IL-33 treatment led to even a significantly higher expansion of infant ST2^+^ Treg cell population (**Supplemental file 6: Figure S6**) suggesting that there is no impairment in IL-33 responsiveness in infant mice. These findings prompted us to investigate whether the infant post-septic condition affects the IL-33 and type 2 cytokines production. We, therefore measured type 2 cytokines (IL-10 and IL-4) and IL-33 production in the lung tissue and bronchoalveolar lavage (BAL) from sepsis-surviving mice. Lung was selected since epithelial and endothelial lung cells are the major sources of IL-33 ^23, 50^. Consistent with previously reported studies ^17^, the post-septic condition resulted in a significant increase in lung IL-33 production and expression as well as an increase of type 2 cytokine production in adult mice. Remarkably, no such increase was evident in post-septic infant mice (**Fig. 4C – F**). Moreover, we observed that whereas post-septic adult mice showed an increase in BAL IL-33 production accompanied by the reduction in the IL-33 soluble receptor (sST2), the production of BAL IL-33 was significantly reduced in post-septic infant mice with no changes in sST2 production (**Supplemental file 7: Figure S7A, B**). Pulmonary epithelial cells, especially pulmonary alveolar type II (Lin^−^EpCAM^+^), have been characterized as the main early and late producers of IL-33 ^23, 24^. Hence, we carried out the expression of *Il33* by lung Lin-cells in post-septic mice. Strikingly, we observed only a significant increase of IL33 expression on adult but not infant Lin-cells (**Fig. 4G**). We then sought to determine the expression of *Il33* in alveolar epithelial cells (AECs) isolated from post-septic infant mice. AECs from septic-surviving infant mice showed a reduction in the expression of *Il33* when compared to the Sham group (**Fig. 4H and Supplemental file 7: Figure S7C**). IL-33 production is finely regulated by both molecular and epigenetic events ^51, 52^. Specifically, in a pathological context, IL-33 expression can be regulated by events of acetylation or methylation ^52, 53^. To further understand the underlying mechanisms involved in the impairment of IL-33 production in post-septic infant mice, we assess the total imprinting DNA methylation of lung Lin-cells from sepsis-surviving infant and adult mice. Strikingly, the post-sepsis condition led to an increase in the total methylation signature of infant Lin-cells compared to that of the adult mice (**Fig. 4I**). Importantly, a significant increment in ILC2s frequency, a common downstream IL-33 target population, was only observed in the adult post-septic group but not in the infant post-septic mice (**Fig. 4J and Supplemental file 7: Figure S7D**). Furthermore, after *in vitro* stimulation, no impairment in the ILCs function was verified in infant mice assessed by the intracellular production of IL-13 (**Fig. 4K**). Collectively, these findings suggest that the infant immunosuppression “resistance” might be associated with the impairment of IL-33 production.

**FIGURE 4.**
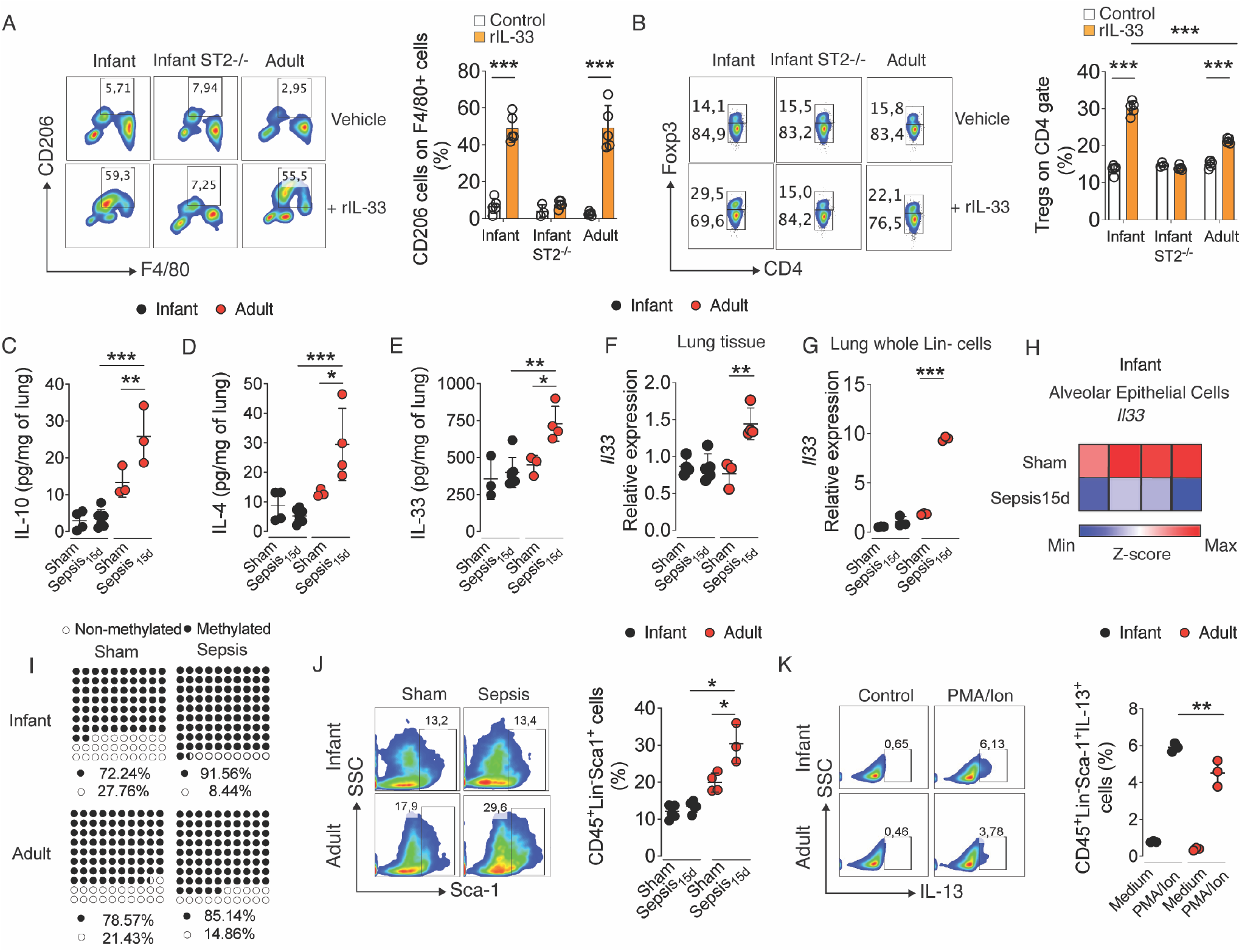
Sepsis does not increase the ILC2/IL-33 axis in sepsis-surviving infant mice. We treated wild type mice (infant and adult) or infant ST2 deficient mice (ST2-/-) with recombinant IL-33 (rIL-33, 0,5μg/kg per 6 days), afterwards, the peritoneal lavage and the spleen were collected, and their cell composition was evaluated by flow cytometry. Representative flow cytometry plots and frequency of **(A)** M2-like macrophages calculated as the percentage of CD206^+^ cells among those F4/80^+^ in the peritoneal exudates and **(B)** spleen Tregs as Foxp3^+^ cells in the gate of those CD4^+^. Infant and adult mice were intraperitoneally injected with 2×10^8^ and 4×10^8^ colony-forming units (CFU) of cecum bacteria, respectively, and after six hours, the ertapenem antibiotic therapy (Abx, 15 mg/kg for infant and 30 mg/kg for adults) was initiated and maintained for 3 days, via intraperitoneal (i.p) twice a day. The following analyses, except the item K, were performed with the 15 days sepsis-surviving (Sepsis_15d_) mice. Concentration in picrogram per mL (pg/mL) of **(C)** IL-10, **(D)** IL-4 and **(E)** IL-33 in the lung homogenates. *Il33* relative expression evaluated by qPCR in **(F)** lung tissue, **(G)** lung whole Lin-cells (CD3ε-Gr-1-CD11b-CD45R/B220-mTer-119) and **H)** alveolar epithelial cells isolated from Sepsis_15d_ infant mice (AECs, CD3ε-Gr-1-CD11b-CD45R/B220-mTer-119-EpCAM^+^). **(I)** total imprinting DNA methylation percentage performed in Lin-cells. **(J)** Representative Flow cytometry plots and frequency in the percentage of lung ILC2s cells (CD45^+^Lin-Sca1^+^ cells). **(K)** Representative Flow cytometry plots and frequency in the percentage of IL13-producing ILC2s cells (CD45^+^Lin^−^Sca-1^+^IL-13^+^) from infant and adult *naïve* mice after phorbol 12-myristate 13-acetate/ionomycin stimulation (PMA/Ion). Data are mean ± SD, n=4-6, representative of two experiments, \**p*<0.05, \*\**p*<0.01, *** *p*<0.001 and **** *p*<0.0001 (**A – G**, and **J, K** one way-ANOVA, Bonferroni’s; **H**, data are Z-score normalized heat map; **I**, data are presented as % of DNA methylated and no methylated).

### Lack of Tregs/IL-33 expansion in sepsis-surviving pediatric patients

Finally, we assessed whether our data from the murine models could be extended to the clinical setting. For that, we investigated the frequency of Tregs in the peripheral blood as well as the serum concentrations of IL-33 in healthy control volunteers and sepsis-surviving adult and pediatric patients. We prospectively included 21 patients (12 children and 9 adults) after hospital discharge in the Emergency Department of a high-complexity hospital. Healthy volunteers (7 healthy children and 12 healthy adults) were included as controls. The baseline demographic and clinical characteristics are summarized in Supplemental file **Table 1**. PRISM and PELOD scores for pediatric patients and SOFA and APACHE II scores for adult patients during their hospitalization were recorded. Whereas sepsis-surviving adults exhibited a significantly elevated level of Tregs cells compared to healthy controls, the sepsis-surviving infant had a low and similar level of Tregs cells as the healthy infants or adults (**Fig. 5A – C)**. Nevertheless, Tregs cells from the surviving infants retained their suppressive activity (**Supplemental file 8: Figure S8**). Moreover, consistent with the mouse model, we did not find significant changes in the plasma levels of IL-33 of post-septic pediatric patients (**Fig. 5D**). These data, therefore demonstrate that the reduced level of IL-33 production in infant mice and pediatric sepsis patients could be responsible for the reduced level of Tregs cells and the resistance of long-term immune suppression that is often observed in adult sepsis individuals.

**FIGURE 5.**
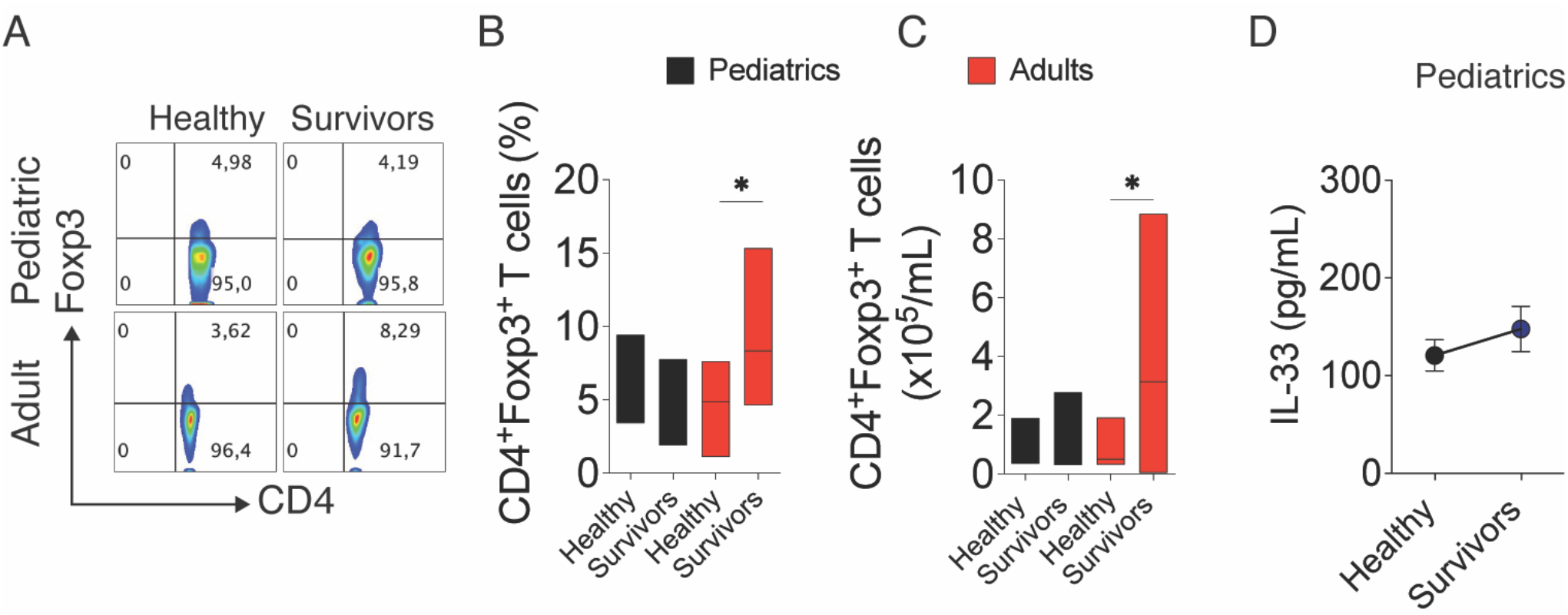
Lack of Tregs/IL-33 expansion in sepsis-surviving pediatric patients. Blood samples were collected from both pediatric and adult sepsis-surviving patients, as well as healthy volunteers, after hospital discharge. **(A)** Representative flow cytometry plots **(B)** frequency in percentage **(C)** and absolute number in 10^5^ cells per milliliter (10^5^/mL) of CD4^+^Foxp3^+^ T cells. **(D)** Serum IL-33 levels in healthy and sepsis-surviving pediatric patients in picogram per mL (pg/mL). Data are mean ± SD, ** *p*<0.01 and **** *p*<0.0001 (Kruskal-Wallis unpaired test).

## Discussion

Several studies have described the post-sepsis immunosuppression syndrome in adults (Cavassani et al., 2010; Hotchkiss et al., 2013; Monneret et al., 2003; Nascimento et al., 2010; Nascimento et al., 2017). However, such immune consequences of sepsis are not significant in children. Concordantly, in comparison to non-survivors, sepsis-surviving pediatric patients did not show leukopenia, an outcome accompanied by reduction of IL-10 expression in monocytes (Hall et al., 2007; Hall et al., 2011). Furthermore, neonate mice with early-onset sepsis did not have an increased frequency for the development of late-onset sepsis immunosuppression (Wynn et al., 2013). Moreover, whereas sepsis-surviving adult patients have up to five times the risk of acquiring a secondary infection after hospital discharge (Cuthbertson et al., 2010), the infant post-septic patients have a low risk of secondary infection (Morrison et al., 2002). The mechanisms for this differential immune suppression between adults and infants recovering from sepsis are largely unknown.

Data reported here demonstrate that, compared to the adults, sepsis-surviving infant mice do not develop post-sepsis immunosuppression. Specifically, post-septic infant mice are resistant to secondary bacterial infections. Notably, we found that this is associated with a failure of Tregs expansion as well as reduced activation of M2-like macrophages/IL-33 axis. Moreover, the post-septic infant condition was associated with an increase in DNA methylation in lung Lin-cells, leading to reduced IL-33 production compared with those from the adult counterparts. Consistent with this observation, treatment of infant mice with exogenous IL-33 led to a higher expansion of the Tregs population and immunosuppression, demonstrating that the decreased IL-33 production in infant mice is essential for their resistance to post-sepsis immunosuppression. The clinical relevance of our findings was supported by the observation that sepsis-surviving pediatric patients exhibit neither systemic Tregs expansion nor increased serum IL-33 levels compared to adult counterparts. Expansion of cord blood Tregs population in neonatal patients with early-onset sepsis (12 days old) has been reported to be inversely correlated with the severity of the disease (Timperi et al., 2016). Likewise, the expansion of the Tregs population has been reported in newborn mice (five to seven days old) 24 h after sepsis induction (Wynn et al., 2007). The discrepancy between these reports and our current finding is likely due to the difference in experimental protocols used. Unlike our experimental system, most of these studies used umbilical cord blood and included acute sepsis patients instead of sepsis-surviving patients. It is noteworthy that Tregs *in vitro* differentiation and CD4 T cells TGF-ß responsiveness are not compromised by age, suggesting that the failure of Tregs expansion *in vivo* does not rely on failure in infants Tregs fitness. Furthermore, the expansion of M2 macrophages in adult post-septic mice was not observed in infants. However, similarly to Tregs, the differentiation of M2 macrophages *in vitro* was not compromised in infant mice. These results suggest that the mechanisms that drive the M2/Treg axis could be upstream of these cells.

Recently, our group demonstrated the role of IL-33 as a key regulator of post-sepsis immunosuppression in adult mice (Nascimento et al., 2017). IL-33 is constitutively produced mainly by epithelial and endothelial cells and acts as an endogenous danger signal, or alarmin, in response to tissue damage (Cayrol and Girard, 2014; Liew et al., 2010; Smith, 2010). IL-33 levels remain elevated in the lung of sepsis-surviving adult mice, leading to the expansion and activation of ILCs populations, which orchestrates a macrophage alternative reprogramming toward an M2-like profile, type 2 cytokines production and expansion of Tregs population (Nascimento et al., 2017). In our study, we observed that the M2-like reprogramming, type 2 cytokines production (such as IL-4), and especially IL-33 production in sepsis-surviving infant mice was not increased, suggesting an unrecognized age-dependent regulation of IL-33 production in a post-septic state. The expression of IL-33 is finely regulated by epigenetic events (Polumuri et al., 2012; Zhang et al., 2014) (Larouche et al., 2018; Zhang et al., 2014) thus we addressed the total DNA methylation imprinting in post-septic infant mice. Compared with adults, lung Lin-cells from post-septic infant mice show a higher total methylation signature, which might be related to the impairment of IL-33 production. The mechanism by which methylation signature might regulate IL-33 production during pediatric sepsis warrants further investigation. Taken together, our findings reveal that post-sepsis immunosuppression is an age-dependent phenomenon. In this context, the differential production of IL-33 has an important implication for the treatment of adult and pediatric post-sepsis immunosuppression.

## Abbreviations

Tregs: Regulatory T cells
BMDM: Bone marrow-derived macrophages
AECs: Alveolar Epithelial Cells
GOT: Glutamic Oxaloacetic Transaminase
ILC2s: Type 2 innate lymphoid cells

## Declarations

### Consent for publication

Not applicable

## Authors’ contributions

Conceived and designed the experiments: DC, CW, WT, VB, FV, DN, JA, BSF and FC. Performed the experiments: DC, CW, WT, VB, MF, FV, DN, DP and MHL. Analyzed the data: DC, CW, FV, JA and FC. Contributed reagents/materials/analysis tools: FC. Wrote/revised the paper: DC, CW, WT, JA, BSF and FC.

## Acknowledgements

We are grateful to Ieda dos Santos, Marco Antônio, Sergio Rosa, Vagner Jesus, Ana Kátia dos Santos and Giuliana Bertozi for technical assistance. We are also grateful to Nathalia Sofia Rosero and Cornelia Rohland for their technical assistance. We also thank the FACS Core Facility of the Medical Faculty at the University of Bonn for providing help, services, and devices funded by the Deutsche Forschungsgemeinschaft (DFG, German Research 764 Foundation, project number 216372545).

## Additional files

**FIGURE S1. Antibiotic-induced recovery from cecal bacteria peritonitis of infant and adult mice and standardization of *P. aeruginosa* sublethal doses used as a second hit in our model**. Experimental setup of our two-hit models with *P. aeruginosa* and B16 tumor cells **(A)**. Infant and adult mice were intraperitoneally injected with 2×10^8^ and 4×10^8^ colony-forming units (CFU) of cecum bacteria, respectively, and after six hours, the ertapenem antibiotic therapy (Abx, 15 mg/kg for infant and 30 mg/kg for adults) was initiated and maintained for 3 days, via intraperitoneal (i.p) twice a day. The survival **(B)** and the variation on the body weight **(C)** expressed in percentage were periodically recorded, as indicated in the figures. In another set of experiments, we evaluated the liver injury by measuring the glutamic-oxalacetic transaminase (GOT) levels expressed in units per milliliter (U/mL) of plasma **(D)**, and the blood bacterial count as the logarithm of CFU per milliliter (Log_10_ CFU/mL) **(E)**, in samples collected from sham animals or septic animals after 6h and 1, 7 and 15 days after infection. Infant **(F)** and adult **(G)** naïve mice were infected with different doses of *P. aeruginosa* (4 × 10^6^, 2 × 10^6^ and 8 × 10^5^ CFUs/mice, intranasally), and the survival was recorded for 10 days and expressed as a percentage. Data are mean ± SD, n=6-8 per group and are representative of 2-3 independent experiments. * *p*<0.5 (*t-*test).

**FIGURE S2. Sepsis-surviving infant mice do not develop CD4 T cells proliferation impairment**. Infant and adult mice were intraperitoneally injected with 2×10^8^ and 4×10^8^ colony-forming units (CFU) of cecum bacteria, respectively, and after six hours, the ertapenem antibiotic therapy (Abx, 15 mg/kg for infant and 30 mg/kg for adults) was initiated and maintained for 3 days, via intraperitoneal (i.p) twice a day. *Ex vivo* proliferation capacity percentage of total spleen CD4 T cells after polyclonal stimulation (anti-εCD3 and anti-CD28, 1μg/mL, respectively) from 15 days sepsis-surviving (Sepsis_15d_) mice. Data are mean ± SD, n=6-8 per group and are representative of 2-3 independent experiments. ** *p*<0.01 (one way-ANOVA, Bonferroni’s).

**FIGURE S3. Sepsis-surviving infant mice do not expand Tregs**. Infant and adult mice were intraperitoneally injected with 2×10^8^ and 4×10^8^ colony-forming units (CFU) of cecum bacteria, respectively, and after six hours, the ertapenem antibiotic therapy (Abx, 15 mg/kg for infant and 30 mg/kg for adults) was initiated and maintained for 3 days, via intraperitoneal (i.p) twice a day. The 15 days sepsis surviving (Sepsis_15d_) mice were then euthanized and the spleen was processed to the flow cytometry analyses **(A)** Representative histogram of and total expression of Foxp3 in CD4^+^ cells as mean fluorescence intensity (MFI). **(B)** Representative flow cytometry plot of Foxp3 expression in CD4^+^ cells. Absolute number of induced **(C)** and natural **(D)** speen-derived T regulatory cells, iTregs and nTregs, based on neuropilin 1 (Nrp1) expression on CD4^+^Foxp3^+^ cells (natural (iTregs: Foxp3^+^Nrp1^−^; nTregs: Foxp3^+^Nrp1^+^). Representative flow cytometry histograms **(E)** and frequency **(F)** in the percentage of spleen Ki67^+^ cells on CD4^+^Foxp3^+^ and CD4^+^ gates from Sepsis_15d_ infant mice. Data are mean ± SD, n=6-8 per group and are representative of 2-3 independent experiments. * *p*<0.05 ** *p*<0.01 and *** *p*<0.001 (**A, C** and **D**, one way-ANOVA, Bonferroni’; **F**, *t-*test).

**FIGURE S4. Tregs *in vitro* differentiation is independent of age. (A)** *In vitro* Tregs differentiation from adult and infant DEREG mice (Foxp3-DTR/EGFP) after stimulation with TGF-β1 (1 and 3 ng/mL) for 72 h, 37°C 5% CO_2_. **(A)** Representative flow cytometry plots and frequency in the percentage of CD4^+^Foxp3eGFP^+^ T cells (Tregs). **(B)** CD4^+^Foxp3eGFP^−^ T cells from infant and adult mice were differentiated *in vitro* and the expression of Tregs hallmark genes were assessed by qPCR. **(C)** Representative flow cytometry plots and frequency of CD25 negative ^(^CD25), CD25 intermediate (CD25^int^) and CD25 high (CD25^hi^) on CD4^+^ T cells after polyclonal stimulation (anti-εCD3 and anti-CD28, 1μg/mL). **(D)** Representative *immunoblots* of p-Smad2/3, Smad2/3, pCREB and CREB on infant CD4^+^Foxp3eGFP^−^ T cells after TGF-β1 stimulation (30 ng/mL). Data are mean ± SD, n=6-8 per group and are representative of 2-3 independent experiments.

**FIGURE S5. Bone marrow-derived macrophages from infant and adult mice are similarly able to polarize to M2-like macrophages**. Bone marrow-derived macrophages (BMDMs) obtained from infant and adult mice were polarized in the presence of IL-4 (10ng/mL) for 48 h. **(A)** Representative histogram and frequency in the percentage of CD206 on F4/80^+^ macrophages. **(B)** M2-like macrophages hallmark genes after *in vitro* polarization in the time point indicated assessed by qPCR. **(C)** CCL22 and IGF-1 levels were expressed in picograms per mL (pg/mL) in the supernatant of the macrophages culture 48 h after stimulation with IL-4. Data are mean ± SD, n=4-6, representative of two experiments, \**p*<0.05, \*\**p*<0.01 and *** *p*<0.001 (one way-ANOVA, Bonferroni’s).

**FIGURE S6. Recombinant IL-33 boosted the expansion of ST2**^**+**^**Tregs**. We treated wild type mice (infant and adult) or infant ST2 deficient mice (ST2-/-) with recombinant IL-33 (rIL-33, 0,5μg/kg per 6 days) and the spleens were collected, and their cell composition was evaluated by flow cytometry. Frequency and representative flow cytometry plots spleen Tregs as Foxp3^+^ cells in the gate of those CD4^+^. Percentage and representative flow cytometry plots of spleen Tregs ST2^+^ (Foxp3^+^ST2^+^ cells). Data are mean ± SD, n=3-5 per group. *p<0. (Two way-ANOVA, Tukey).

**FIGURE S7. Sepsis does not increase the ILC2/IL-33 axis in sepsis-surviving infant mice**. Infant and adult mice were intraperitoneally injected with 2×10^8^ and 4×10^8^ colony-forming units (CFU) of cecum bacteria, respectively, and after six hours, the ertapenem antibiotic therapy (Abx, 15 mg/kg for infant and 30 mg/kg for adults) was initiated and maintained for 3 days, via intraperitoneal (i.p) twice a day. The 15 days sepsis surviving (Sepsis_15d_) mice were then euthanized and the bronchoalveolar lavage (BAL) and the lung tissue were collected for further analysis. **(A)** IL-33 and **(B)** sST2 levels in picogram per mL (pg/mL) in the BAL. **(C)** Sorting strategy flow cytometry plots of lung alveolar epithelial cells (AECs). **(D)** Absolute number of lungs ILC2s cells (CD45^+^Lin^−^Sca1^+^ cells) expressed in cells × 10^5^ (×10^5^), mean fluorescence intensity × 10^3^ (MFI × 10^3^) and Sca1 expression in percentage on CD45^+^Lin^−^ cells. Data are mean ± SD, n=4-6, representative of two experiments, \**p*<0.05 (one way-ANOVA, Bonferroni’s).

**FIGURE S8. Tregs suppression activity is preserved in sepsis-surviving pediatric patients**. Blood samples were collected from sepsis-surviving pediatric patients, and healthy volunteers, after hospital discharge. Representative histogram of T effector cells (CD4^+^CD25^−^CFSE^+^) proliferation in the presence of a serially diluted concentration of Tregs 72 h after polyclonal stimulation (1μg/mL).

**TABLE S1**. Demographic and clinical characteristics of the sepsis-surviving patients. SOFA, Sequential Organ Failure Assessment. APACHE, Acute Physiology and Chronic Health Evaluation. PRISM, Pediatric Risk of Mortality. PELOD, Pediatric Logistic Organ Dysfunction. PRISM, Pediatric Risk of Mortality.

