## Supplemental files for "IL-33 regulates age-dependency of long-term immune dysfunction induced by sepsis"

[1]Center of Research in Inflammatory Diseases (CRID), Ribeirão Preto Medical School, Departments of [2]Biochemistry and Immunology and [3]Pharmacology, University of São Paulo, Ribeirão Preto Brazil. [4]School of Dentistry, Alfenas Federal University, Alfenas, Brazil. [5]Pediatrics and [6]Pathology, University of São Paulo, Ribeirão Preto, Brazil. [7]Institute of Innate Immunity, Medical Faculty, University of Bonn, 53127 Bonn, NRW, Germany.

***Correspondence:** Fernando Q Cunha; Center of Research in Inflammatory Diseases (CRID), Ribeirão Preto Medical School, Rua das Paineiras, casa 3; 14049-900, Ribeirao Preto, SP, Brazil. Tel: +55 16 33153324;

**TABLE S1.** Demographic and clinical characteristics of the sepsis-surviving patients. SOFA, Sequential Organ Failure Assessment. APACHE, Acute Physiology and Chronic Health Evaluation. PRISM, Pediatric Risk of Mortality. PELOD, Pediatric Logistic Organ Dysfunction. PRISM, Pediatric Risk of Mortality.

### Figures

Figure S1

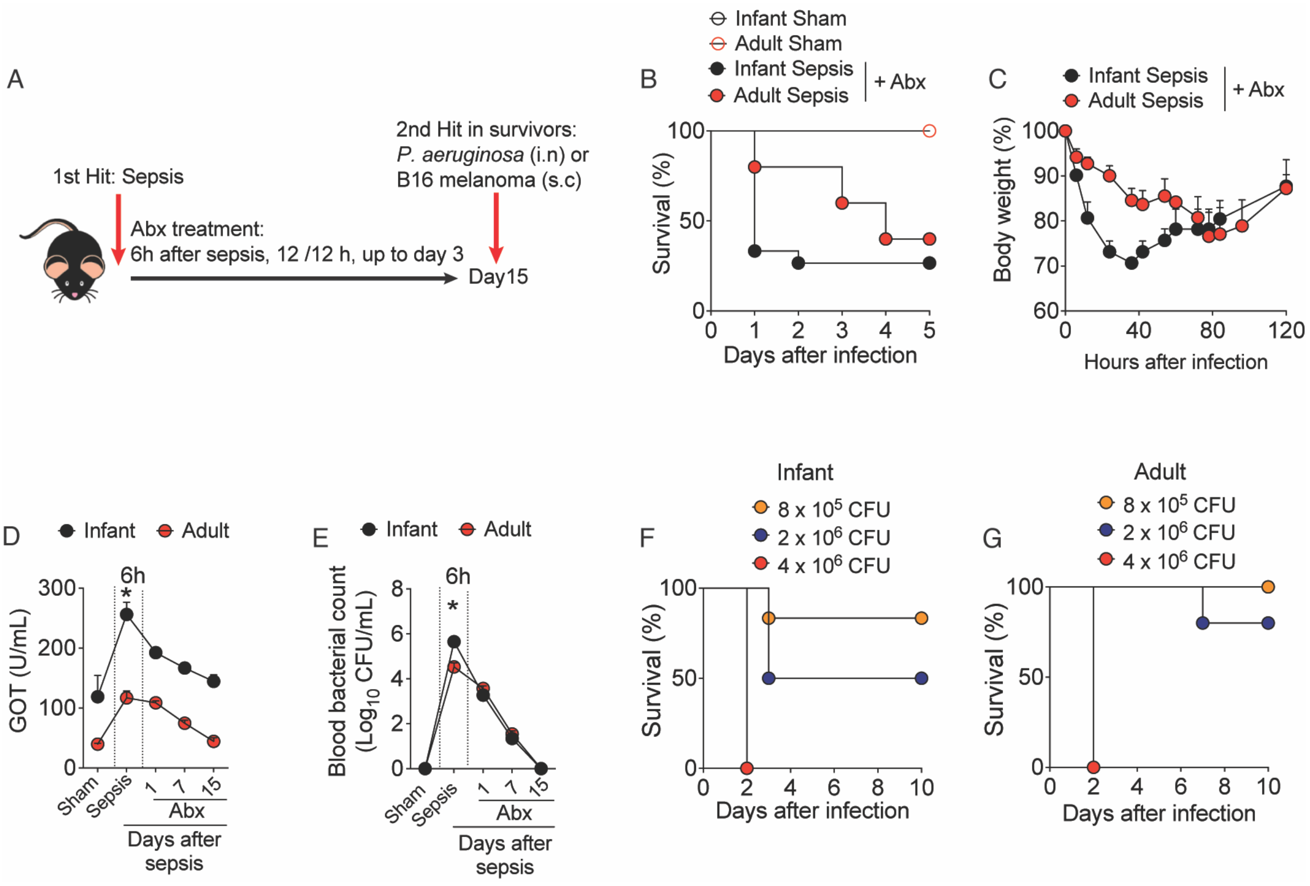

Figure S2

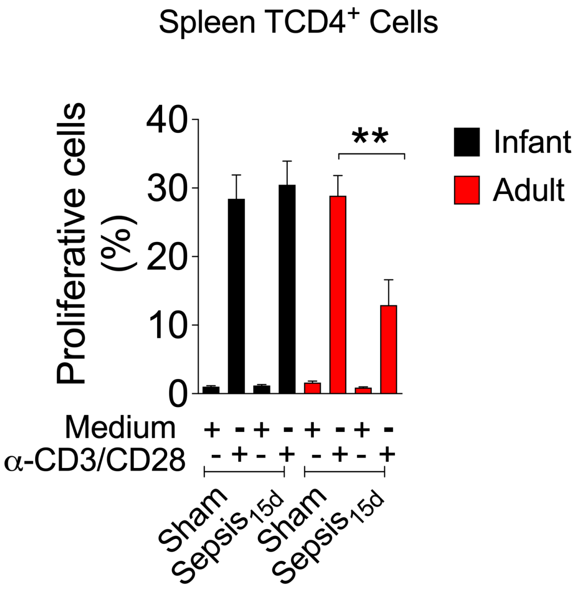
Spleen TCD4+ Cells
Proliferative cells (%)
40
30
20
10
0
**
Infant
Adult
Medium + - + - + - + -
α-CD3/CD28 - + - + - + - +
Sham
Sepsis15d
Sham
Sepsis15d

Figure S3

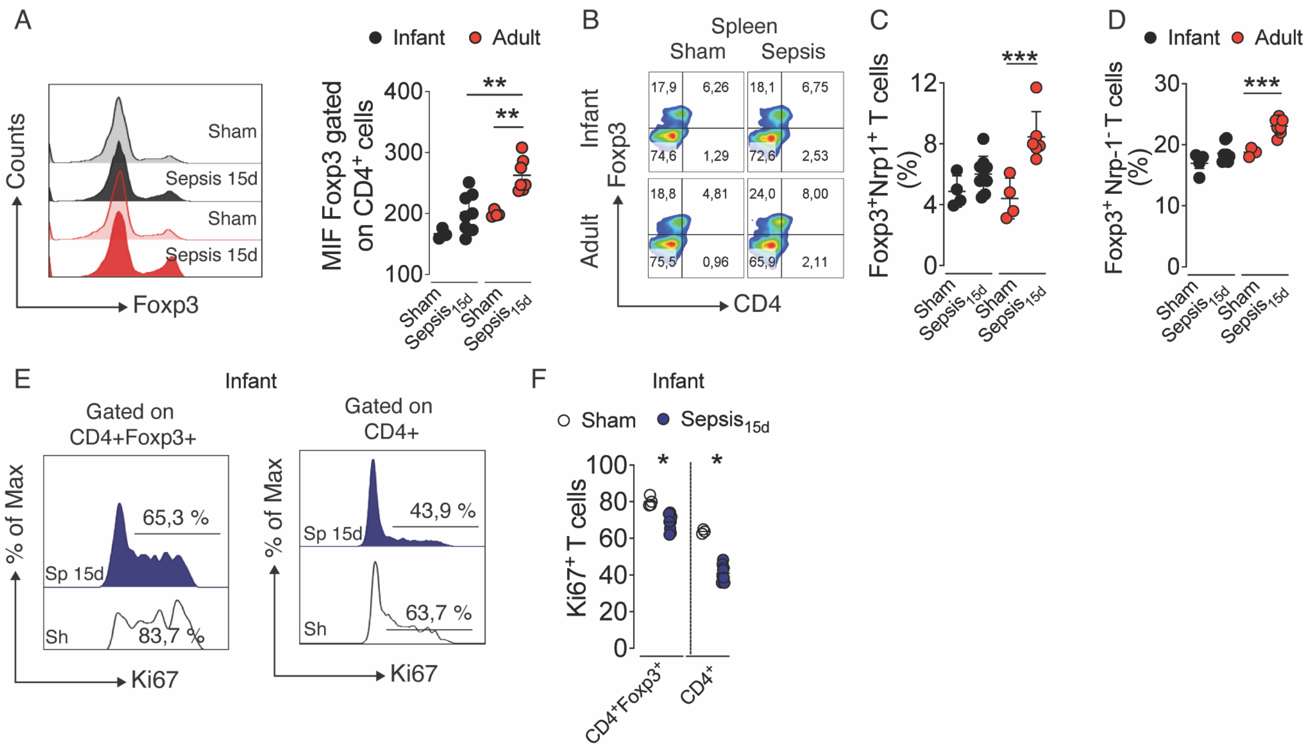
A
Infant
Adult
Counts
Sham
Sepsis 15d
Sham
Sepsis 15d
Foxp3
MIF Foxp3 gated on CD4+ cells
400
300
200
100
**
**
Sham
Sepsis15d
Sham
Sepsis15d
B
Spleen
Sham
Sepsis
Infant
Adult
Foxp3
CD4
17,9
6,26
74,6
1,29
18,1
6,75
72,6
2,53
18,8
4,81
75,5
0,96
24,0
8,00
65,9
2,11
C
Foxp3+Nrp1+ T cells (%)
12
8
4
0
***
Sham
Sepsis15d
Sham
Sepsis15d
D
Infant
Adult
Foxp3+ Nrp-1- T cells (%)
30
20
10
0
***
Sham
Sepsis15d
Sham
Sepsis15d
E
Infant
Gated on CD4+Foxp3+
Gated on CD4+
% of Max
65,3 %
Sp 15d
Sh
83,7 %
Ki67
43,9 %
Sp 15d
Sh
63,7 %
Ki67
F
Infant
Sham
Sepsis15d
Ki67+ T cells
100
80
60
40
20
0
*
*
CD4+Foxp3+
CD4+

Figure S4

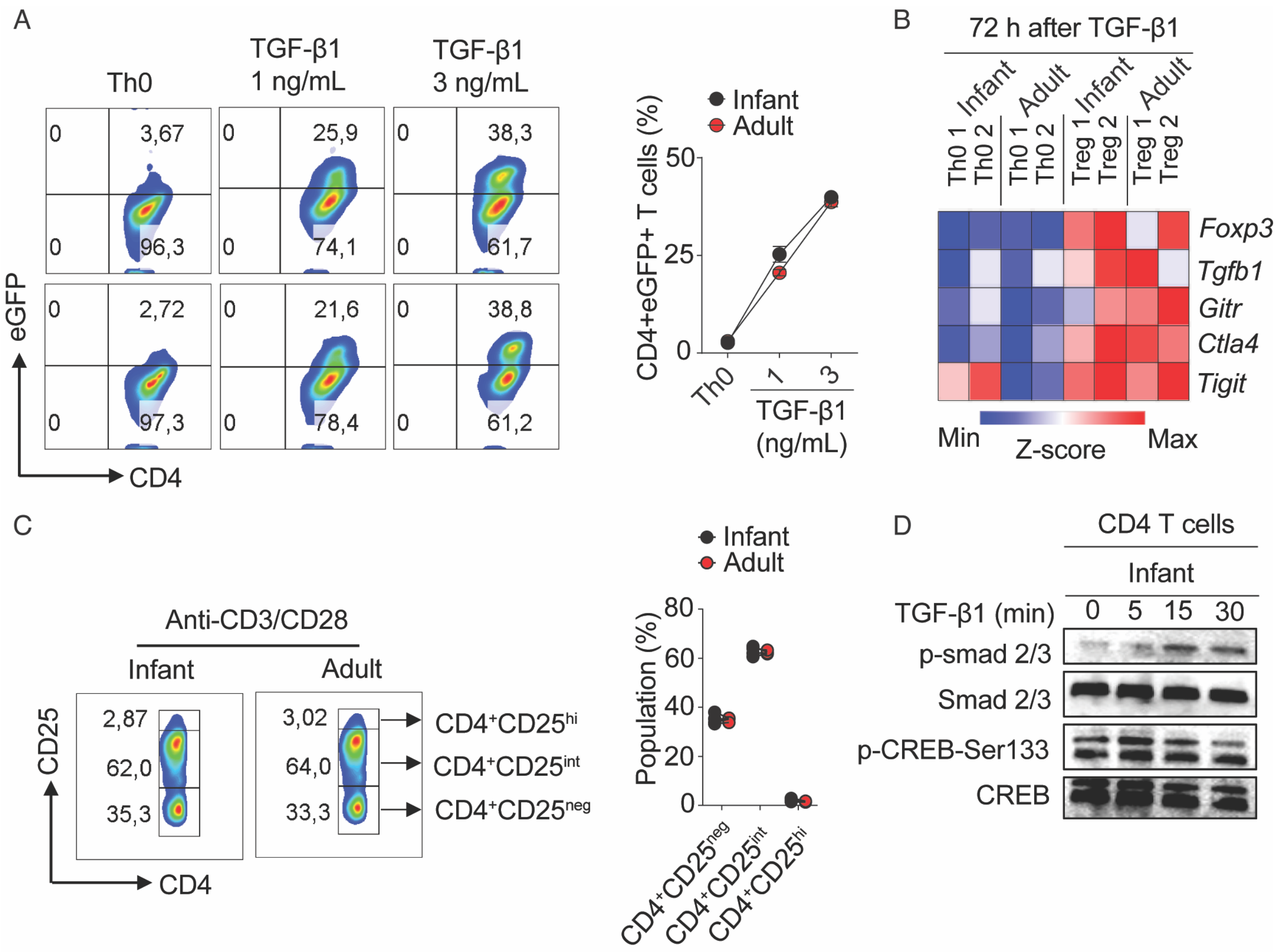
A
Th0
TGF-β1
1 ng/mL
TGF-β1
3 ng/mL
3,67
96,3
25,9
74,1
38,3
61,7
2,72
97,3
21,6
78,4
38,8
61,2
eGFP
CD4
Infant
Adult
CD4+eGFP+ T cells (%)
50
25
0
Th0
1
3
TGF-β1
(ng/mL)
B
72 h after TGF-β1
Infant
Adult
Infant
Adult
Th0 1
Th0 2
Th0 1
Th0 2
Treg 1
Treg 2
Treg 1
Treg 2
Foxp3
Tgfb1
Gitr
Ctla4
Tigit
Min
Z-score
Max
C
Anti-CD3/CD28
Infant
Adult
2,87
62,0
35,3
3,02
64,0
33,3
CD25
CD4
CD4+CD25hi
CD4+CD25int
CD4+CD25neg
Infant
Adult
Population (%)
80
60
40
20
0
CD4+CD25neg
CD4+CD25int
CD4+CD25hi
D
CD4 T cells
Infant
TGF-β1 (min)
0
5
15
30
p-smad 2/3
Smad 2/3
p-CREB-Ser133
CREB

Figure S5

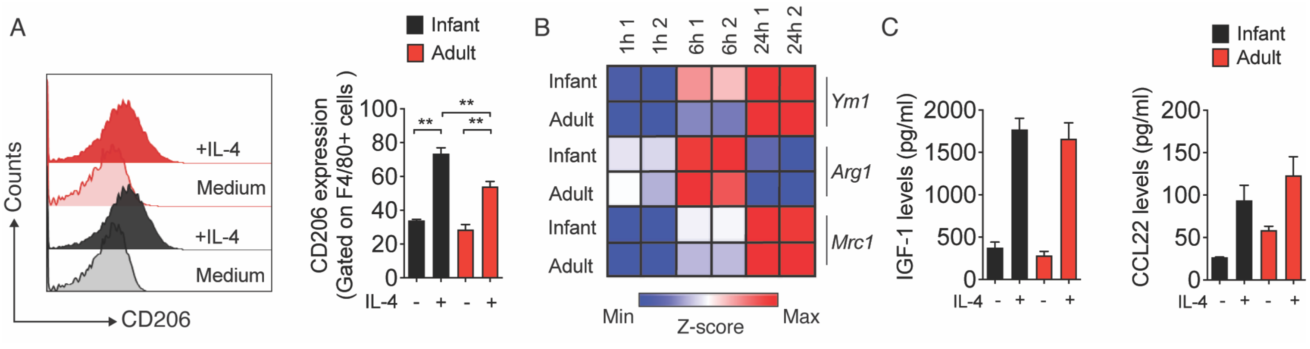
A
Infant
Adult
Counts
+IL-4
Medium
+IL-4
Medium
CD206
CD206 expression
(Gated on F4/80+ cells )
100
80
60
40
20
0
**
**
**
IL-4 - + - +
B
1h 1
1h 2
6h 1
6h 2
24h 1
24h 2
Infant
Adult
Infant
Adult
Infant
Adult
Ym1
Arg1
Mrc1
Min
Z-score
Max
C
Infant
Adult
IGF-1 levels (pg/ml)
2000
1500
1000
500
0
IL-4 - + - +
CCL22 levels (pg/ml)
200
150
100
50
0
IL-4 - + - +

Figure S6

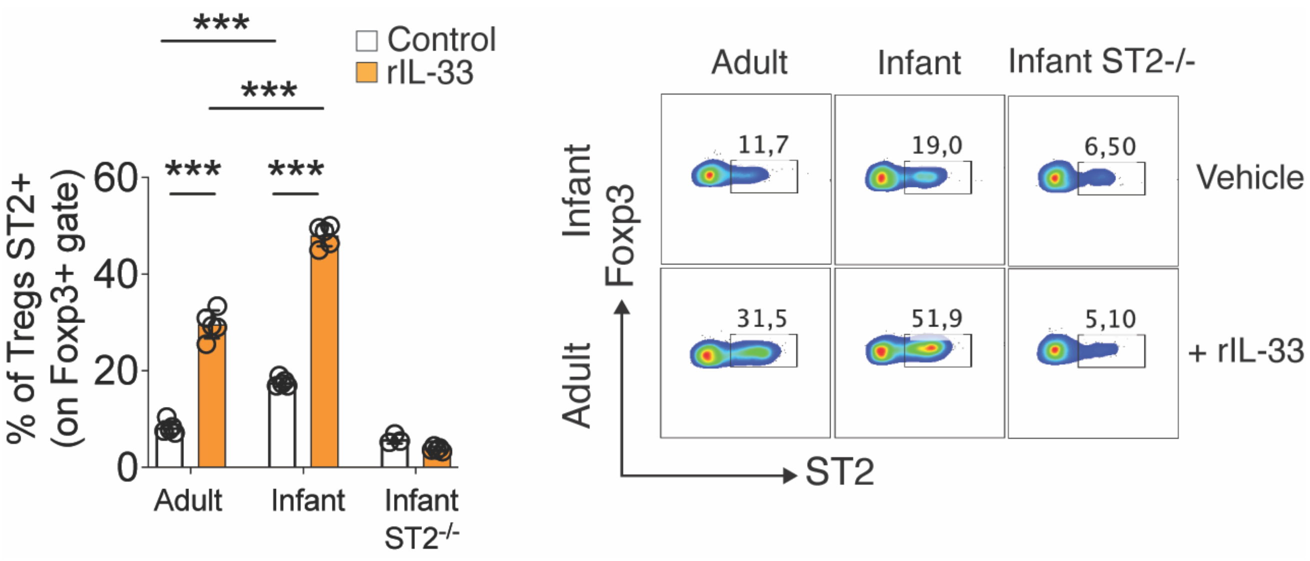

***
***
Control
rIL-33
*** ***
% of Tregs ST2+
(on Foxp3+ gate)
60
40
20
0
Adult
Infant
Infant ST2-/-
Adult
Infant
Infant ST2-/-
11,7
19,0
6,50
Vehicle
31,5
51,9
5,10
+ rIL-33
Foxp3
ST2

Figure S7

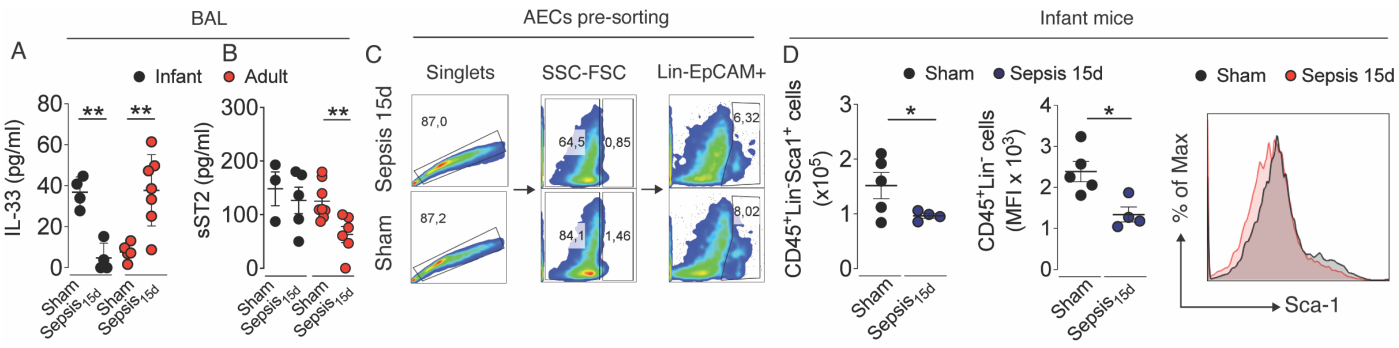
BAL
A
B
Infant
Adult
IL-33 (pg/ml)
sST2 (pg/ml)
Sham
Sepsis15d
AECs pre-sorting
C
Singlets
SSC-FSC
Lin-EpCAM+
Sepsis 15d
Sham
87,0
87,2
64,5
0,85
84,1
1,46
6,32
8,02
Infant mice
D
Sham
Sepsis 15d
CD45+Lin-Sca1+ cells (x10^5)
CD45+Lin- cells (MFI x 10^3)
% of Max
Sca-1

Figure S8

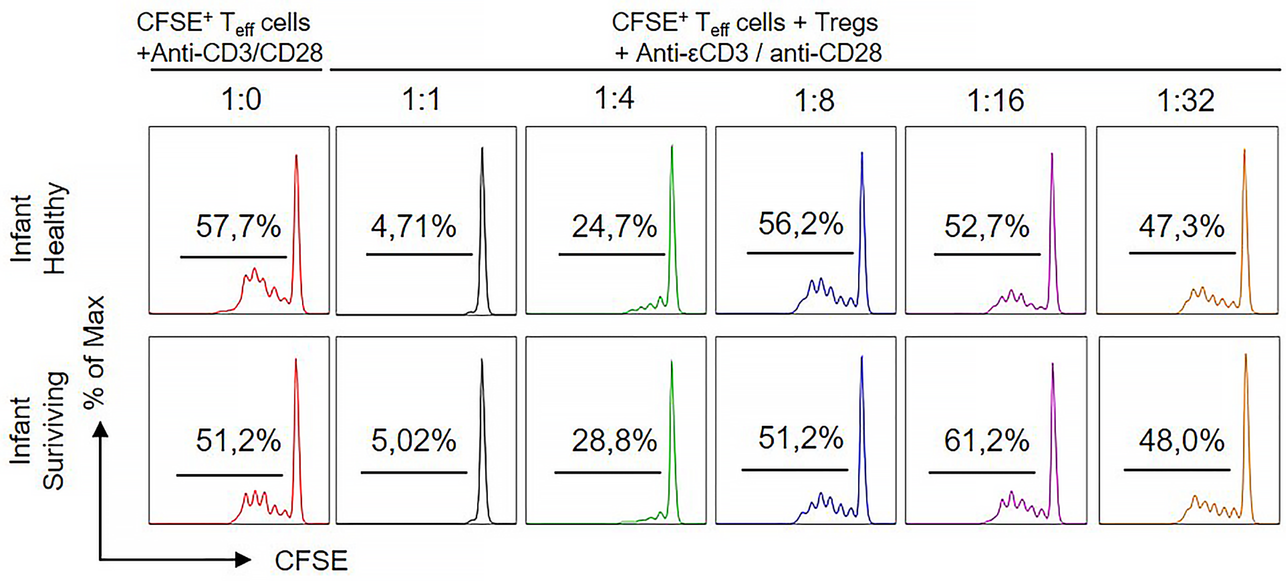
CFSE+ Teff cells
+Anti-CD3/CD28
CFSE+ Teff cells + Tregs
+ Anti-εCD3 / anti-CD28
1:0
1:1
1:4
1:8
1:16
1:32
Infant
Healthy
57,7%
4,71%
24,7%
56,2%
52,7%
47,3%
Infant
Suriviving
51,2%
5,02%
28,8%
51,2%
61,2%
48,0%
% of Max
CFSE

**TABLE 1.** Baseline demographic and clinical characteristics of the sepsis-surviving patients.

| Characteristics | Pediatric Patients (n=12) | Adult Patients (n=9) |
|---|---|---|
| **Age (yrs) – mean (SEM)** | 4.69 (±4.36) | 53.43 (±17.41) |
| **Female – n (%)** | 3 (25%) | 5 (55%) |
| **APACHE II – mean (SEM)** | N/A | 21.33 (±9.89) |
| **SOFA – mean (SEM)** | N/A | 10 (±4.06) |
| **PRISM – mean (SEM)** | 5.66 (±2.83) | N/A |
| **PELOD – mean (SEM)** | 5.83 (±4.72) | N/A |

*N/A – Non-applied

SOFA, Sequential Organ Failure Assessment. APACHE, Acute Physiology and Chronic Health Evaluation. PRISM, Pediatric Risk of Mortality. PELOD, Pediatric Logistic Organ Dysfunction. PRISM, Pediatric Risk of Mortality.
